# Morphological and transcriptomic renal characteristics of dioxin-induced fetal obstructive uropathy

**DOI:** 10.64898/2026.11.23.768580

**Authors:** Mayke A.C. ten Hoor, Robert van Leeuwen, Karamjit Singh Dolt, Peter Hohenstein, Jaap Mulder

**Affiliations:** Department of Human Genetics, Leiden University Medical Center, Leiden, The Netherlands; Division of Nephrology, Department of Pediatrics, Willem-Alexander Children’s Hospital, Leiden University Medical Center, Leiden, The Netherlands; Central Animal Facility, Leiden University Medical Center, Leiden, The Netherlands; Transgenic Facility Leiden, Central Animal Facility, Leiden University Medical Center, Leiden, The Netherlands; Division of Nephrology, Department of Pediatrics, Sophia Children’s Hospital, Erasmus Medical Center, Rotterdam, The Netherlands

**Keywords:** Dioxin, COU, Mouse model, TCDD

## Abstract

Congenital obstructive uropathy (COU) is an important cause of childhood kidney failure, yet understanding its variable renal outcomes and genetic modifiers is limited by the lack of genotype-independent models of *in utero* obstruction. Here, we characterized renal morphology and gene expression at embryonic day (E)18.5 in a mouse model of COU induced by exposure to 2,3,7,8-tetrachlorodibenzo-*p*-dioxin (dioxin) at E10.5. Dioxin exposure reproducibly induced bilateral hydroureteronephrosis with selective medullary flattening with relatively preserved renal architecture. Transcriptomic analysis revealed altered developmental, cell cycle, metabolic, innate immune, and toxin response programs, alongside changes in cell-type-associated signatures. However, immunohistochemistry showed no differences in the abundance of examined stromal and proximal tubular populations, suggesting altered cellular states rather than changes in cellular composition. Together with morphology consistent with early obstructive responses and the absence of prominent injury markers, these findings support its use as a non-genetic model to investigate how congenital obstruction perturbs renal development.

## Introduction

Congenital obstructive uropathy (COU) comprises developmental abnormalities that impair urinary outflow, including ureteropelvic junction obstruction, ureterovesical junction obstruction, ureteroceles, and lower urinary tract obstruction such as posterior urethral valves. Obstructive uropathies occur in approximately 1:2000 pregnancies and encompass a broad phenotypic spectrum with highly variable clinical outcomes [1–3]. As COU represents an important cause of kidney failure in children, understanding the mechanisms that determine the severity and progression of kidney injury is essential for improving prognostication and treatment [3–5].

Urinary tract obstruction can increase upstream pressure, leading to hydronephrosis, defined as dilation of the renal collecting system, which may subsequently compress the developing renal parenchyma. As obstruction occurs during active nephrogenesis, hydronephrosis can be accompanied by renal hypoplasia, characterized by reduced kidney growth and nephron mass, and dysplasia, resulting from abnormal renal differentiation [6]. These developmental abnormalities may compromise nephron maturation and final nephron endowment, with obstruction occurring earlier in gestation generally associated with more severe outcomes [7–9]. In parallel, increased pressure initiates interconnected processes involving glomerular and tubular injury, oxidative stress, inflammation, cell death, and fibrogenic signaling, while persistent obstruction may ultimately lead to tubular atrophy, interstitial fibrosis, and nephron loss [3, 10, 11].

The effects of urinary tract obstruction on kidney development, subsequent renal injury, and the contribution of genetic factors have been investigated using a range of experimental models, each capturing different aspects of COU. Surgically induced unilateral ureteral obstruction (UUO) can be performed in utero in large animals, including fetal sheep, reproducing developmental consequences such as impaired nephrogenesis and renal dysplasia but requires invasive intrauterine procedures [9, 12]. In rodents, UUO is widely used in adults to study tubular injury, inflammation, and fibrosis, whereas neonatal UUO additionally models disruption of ongoing nephrogenesis but is technically more challenging [13, 14]. Mouse models that develop urinary tract obstruction in utero, such as the megabladder mouse, rely on underlying genetic mutations to induce obstruction [15, 16]. Therefore, there remains a need for a reproducible mouse model in which obstruction arises during embryonic development independently of the experimental genotype, allowing the study of obstruction-induced renal abnormalities and genetic factors that modify them.

A potentially suitable mouse model is fetal obstructive uropathy induced by 2,3,7,8-tetrachlorodibenzo-*p*-dioxin (hereafter, dioxin), a developmental toxicant that acts through activation of the aryl hydrocarbon receptor (AhR) [17–19]. Following in utero exposure, ureteral epithelial hyperplasia causes luminal obstruction by embryonic day (E)15.5, followed by pronounced hydroureteronephrosis by E16.5-E17.5 [20–23]. Importantly, as in human COU, hydroureteronephrosis results from obstruction of urinary outflow, supporting the relevance of this model.

Despite decades of research on dioxin teratogenicity, the broader renal transcriptional features of dioxin-induced COU remain poorly characterized. Here, we combine morphological assessment at E18.5 with transcriptomic profiling to identify biological processes altered in the affected fetal kidney and solidify the utility of this model for studying COU.

## Results

### Dioxin reproducibly induces bilateral hydroureteronephrosis in fetuses

Timed matings were performed, and potentially pregnant females received either control or dioxin gavage at E10.5 (Fig. 1A). Female body weight and behavior were monitored at multiple timepoints to assess the effects of treatment. Dioxin did not affect body weight trajectories, expressed as percentage of weight change from the day post-mating, in either pregnant or non-pregnant females (three-way ANOVA, p = 0.7808). No behavioral abnormalities or signs of discomfort were observed in any group. At E18.5, the numbers of implantations, live fetuses, and resorptions were assessed. Neither the number of implantations per litter (unpaired t-test, p = 0.391) nor the proportion of live fetuses versus resorptions (binomial GLM, p = 0.422) differed between control and dioxin-exposed litters (Fig. 1C). Fetuses were subsequently assessed for urinary tract abnormalities (Fig. 1D). All dioxin-exposed fetuses displayed bilateral hydroureteronephrosis, characterized by tortuous, dilated ureters and dilation of the renal pelvis. In stark contrast, only one control fetus displayed unilateral hydroureteronephrosis and one unilateral renal hypoplasia. The case in the control group may reflect the sporadic occurrence of hydroureteronephrosis in mice, as previously reported both in control fetuses from dioxin exposure studies and during necropsy of several other inbred mouse strains [24, 25]. Taken together, these findings suggest that a single dose of 15 µg/kg dioxin did not affect maternal body weight, behavior, or fetal survival, while reproducibly inducing bilateral hydroureteronephrosis, consistent with previous studies [26–28].

**Figure 1.**
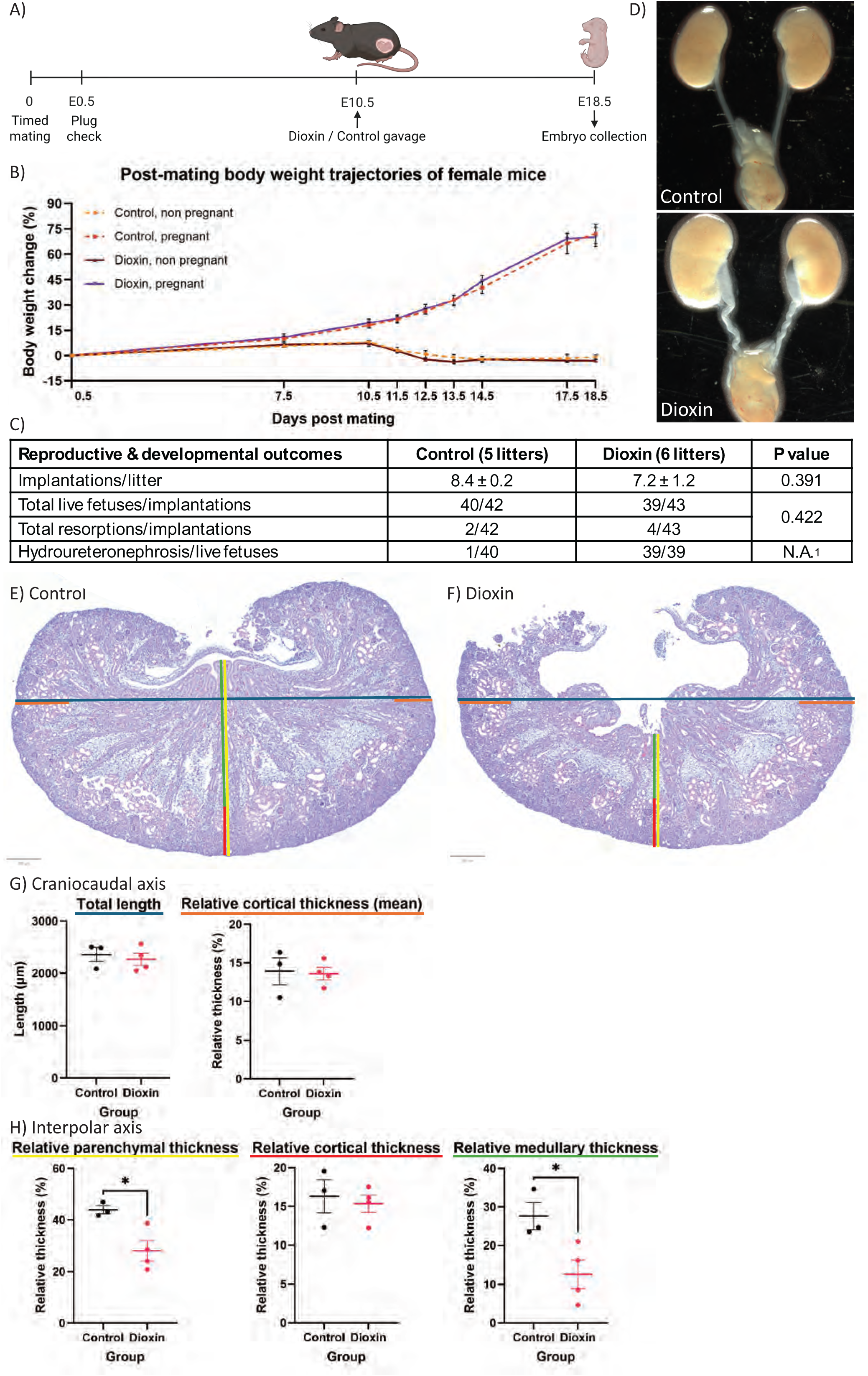
| Pregnancy outcomes and morphology of E18.5 control and dioxin-exposed kidneys. A) Summary of experimental timeline. Created in BioRender. ten Hoor, M. (2026) https://BioRender.com/35vtlof B) Post-mating body weight trajectories, expressed as percentage change from body weight at 0.5 days post mating. C) Pregnancy outcomes in control and dioxin-exposed litters, number of implantations per litter as mean ± SEM, total counts of live fetuses and resorptions per total implantations, and hydroureteronephrosis in live fetuses per litter. ^1^ not possible to perform binomial GLM due to near-complete separation D) Representative gross morphology of the urinary tract from a control embryo with normal kidneys and ureters, and from a dioxin-exposed embryo with hydroureteronephrosis. H&E staining of E) dioxin-exposed and F) control kidneys. The lines indicate the measurements performed, with line colors corresponding to the measurements shown in G) total kidney length and relative cortical thickness (mean) along the craniocaudal axis, and H) relative parenchymal, medullary, and cortical thickness compared to craniocaudal length along the interpolar axis. Data are presented as mean ± S.E.M. (control, n = 3; dioxin, n = 4). * p<0.05.

### Dioxin-induced hydronephrosis is associated with renal pelvic dilation and medullary flattening

The effects of dioxin exposure on E18.5 fetal renal morphology were further assessed by hematoxylin and eosin (H&E) staining. Control kidneys displayed a normal morphology (Fig. 1F), whereas dioxin-exposed kidneys displayed a markedly dilated renal pelvis (Fig. 1G), consistent with hydroureteronephrosis. Morphometric analysis revealed no difference between treatment groups in craniocaudal kidney length (Welch’s t-test, p = 0.6450; Fig. 1H) or mean cortical thickness at the cranial and caudal poles, expressed relative to craniocaudal length (Welch’s t-test, p = 0.8807). At the interpolar axis, relative parenchymal thickness was significantly reduced in dioxin-exposed kidneys (Welch’s t-test, p = 0.0197; Fig. 1I). This reduction was primarily attributable to decreased relative medullary thickness (Welch’s t-test, p = 0.0332), whereas relative cortical thickness was unchanged (Welch’s t-test, p = 0.7177). The reduced medullary thickness was unlikely to result from sectioning plane, as pelvic dilation persisted across sections (data not shown). Collectively, these morphometric measurements indicate that dioxin-induced hydroureteronephrosis is associated with selective flattening of the interpolar medulla, whereas kidney length and cortical thickness remain preserved. This pattern is consistent with the compression of the nephrogenic blastema previously described in the dioxin model [23] and resembles observations in human fetuses with early urinary tract obstruction, where medullary flattening occurs despite otherwise preserved renal architecture [6]. A similar phenotype was observed in the megabladder mouse, in which hydroureteronephrosis at E17.5 was not accompanied by detectable changes in a selected panel of pathogenic markers [29]. Together, these findings support the interpretation that the E18.5 dioxin-induced phenotype reflects an early stage of obstructive nephropathy, characterized by flattening of the medulla before more widespread morphological evidence of renal injury becomes apparent.

### Transcriptional programs in developing kidneys are altered by dioxin exposure

The transcriptional response of the fetal kidney to dioxin exposure was characterized by 3’-end bulk RNA sequencing of whole-kidney lysates collected at E18.5 (n=4 samples per group from different litters). Following mouse reference genome alignment and filtering to remove low expression genes, a total of 11,228 genes were identified for further analysis. Principal component analysis (PCA) showed tight clustering of controls, whereas dioxin-exposed samples were more dispersed, with D4 positioned close to the control group (Fig. 2A). Differential expression analysis using an absolute fold-change cut-off of 1.5, a nominal p-value <0.05, and a non-missing FDR-adjusted p-value identified 567 differentially expressed genes (DEGs) following dioxin exposure, with more genes upregulated than downregulated (412 versus 155; Fig. 2B).Among the most strongly upregulated genes were Cyp1a1, a well-established target of dioxin-induced AhR activation, and S100a9, which has previously been reported to increase in thymocytes following dioxin exposure [24, 30, 31]. Consistent with the PCA, hierarchical clustering of all DEGs placed D4 among the controls, whereas the remaining dioxin-exposed samples formed a distinct expression pattern (Fig. 2C). Although all dioxin-exposed fetuses showed visible hydroureteronephrosis, kidneys used for RNA sequencing were immediately snap-frozen upon harvesting, which precluded histological assessment of obstruction severity in those individual kidneys. Hence, differences in lesion severity and dioxin responsiveness, may have contributed to the observed interindividual transcriptional variability.

**Figure 2.**
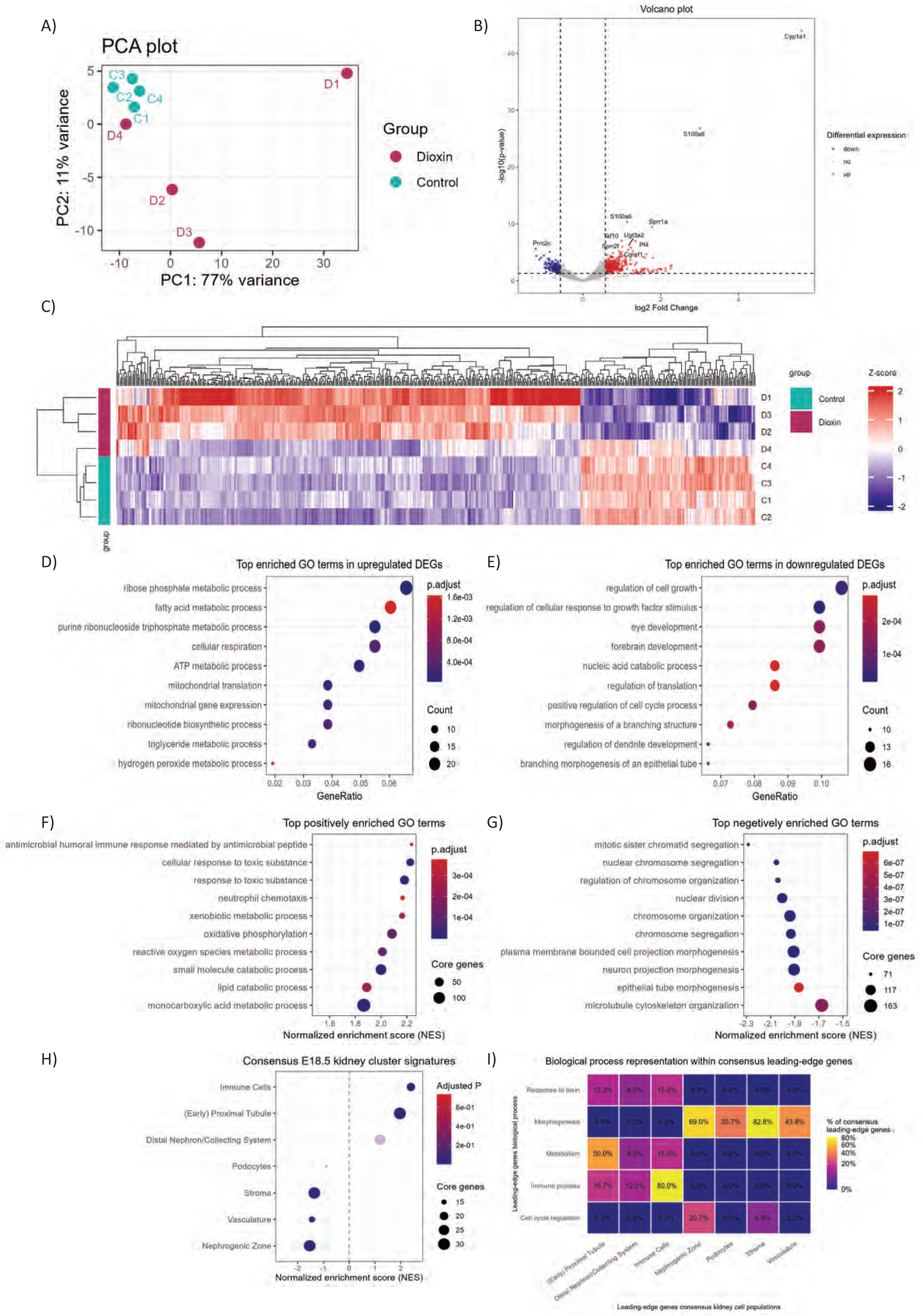
| Transcriptomic effects of dioxin exposure. A) PCA of all samples. B) Volcano plot of DEGs using *p* < 0.05 and |log2FC| > 0.58; the 10 most significant genes are labelled, and genes with missing adjusted *p*-values were excluded. C) Z-score-normalized heatmap of the 50 DEGs based on the lowest adjusted *p*-values. D-E) Dotplot of top 10 simplified GO-BP terms for upregulated and downregulated DEGs, respectively. F-G) Dotplot of top 10 simplified GO-BP terms positively and negatively enriched by GSEA, respectively. H) Dot plot of all consensus kidney cell population signatures tested by GSEA. Dot opacity indicates statistical significance, with faded dots representing non-significant terms (FDR ≥ 0.05). I) Heatmap showing the percentage overlap between leading-edge genes from major enriched biological processes and consensus kidney cell-population signatures.

To assess the biological relevance of the DEGs, gene ontology (GO) enrichment analysis was performed for the biological process category (GO-BP), yielding 43 enriched terms among upregulated and 166 among downregulated DEGs (Supplementary file 1; Tab GOBP – upregulated DEGs / GOBP-downregulated DEGs). Upregulated DEGs were predominantly enriched for metabolic processes, with top terms as “mitochondrial translation” and “cellular respiration” (Fig. 2D). Consistent with dioxin’s known toxicity, “response to toxic substance” was also enriched among upregulated GO-BP terms. In contrast, downregulated DEGs were enriched for morphogenesis-related terms, including “branching morphogenesis of an epithelial tube” and “regulation of cell growth” (Fig. 2E).

With the aim of capturing coordinated expression changes that cannot be detected by the threshold-based GO analysis, gene set enrichment analysis (GSEA) was conducted on the full ranked gene list (Supplementary file 1; Tab GSEA_GOBP). GSEA confirmed the positive enrichment of toxin-response and metabolic terms identified by GO, including “cellular response to toxic substance” and “oxidative phosphorylation”, respectively, and additionally revealed a broader positive enrichment signature encompassing immune-related processes (Fig. 2F). Moreover, GSEA prominently highlighted downregulation of cell cycle programs, with “nuclear division” among the top terms, alongside morphogenesis-related terms such as “morphogenesis of an epithelium” (Fig. 2G). The combined GO and GSEA outcomes indicate a correlation between dioxin exposure and positive enrichment of toxic substance response, metabolic activity, and immune signaling processes, concurrent with the negative enrichment of renal morphogenesis and cell cycle regulation.

### Mapping of transcriptional changes to kidney cell population signatures

Given the reduced interpolar medullary thickness following dioxin exposure, we assessed cell population-associated transcriptional changes by GSEA using cell population consensus signatures derived from two independent E18.5 mouse kidney single-cell RNA-sequencing datasets (Fig. 2H) [32, 33]. As these signatures comprised cell population-enriched genes, enrichment direction reflected higher or lower expression of the corresponding population signature. Across the consensus cell populations, the immune cell and (early) proximal tubule signatures were significantly increased, while nephrogenic zone, stroma, and vasculature signatures were depleted. Podocyte and distal nephron/collecting system signatures were unchanged. This pattern argues against a simple loss of medullary identity with relative cortical enrichment and instead points to a broader change across multiple developmental cell populations in the hydronephrotic dioxin kidneys.

To link the major enriched biological processes to specific cell populations, we compared leading-edge genes, representing the subset of genes contributing most strongly to each GSEA enrichment signal, from terms related to toxin response, morphogenesis, metabolism, immune processes, and cell cycle regulation with those from the consensus cell-population signatures (Fig. 2I). Focusing on overlaps of ≥50%, metabolic genes strongly overlapped with (early) proximal tubule signatures, immune-process genes with immune-cell signatures, and morphogenesis-related genes with both nephrogenic zone and stromal signatures. These findings associate the metabolic, immune, and morphogenesis-related transcriptional changes to distinct kidney cell population signatures.

### Altered developmental programs are associated with multiple renal compartments

Metanephric kidney development relies on reciprocal interactions among the ureteric bud, cap mesenchyme, renal stroma and developing endothelium. Interconnected signaling pathways coordinate ureteric-bud branching for collecting-system formation, differentiation of cap mesenchyme progenitors into nephrons, and development of the renal vasculature [34]. The GSEA identified negative enrichment of 116 development– and morphogenesis-related processes in the dioxin group. Recurrent leading-edge genes of these terms mapped to several interconnected developmental pathways. These pathways comprised Wnt (*Ctnnb1*, *Wnt5a*, *Sfrp1*, *Sfrp2*), Hedgehog (*Ptch1*, *Smo*, *Gli2*), TGF-β/BMP (*Tg@2*, *Tg@r1*, *Bmp2*, *Smad4*), Slit-Robo (*Slit2*, *Robo2*), and Notch signaling (*Notch1*, *Notch2*, *Hey1*) [34]. Moreover, several transcription factors (*Sall1*, *Pax2*, *Lhx1*, *Hnf1b*, *Gata3*, *Foxd1*, *Pbx1*) with established roles in kidney development were among the leading-edge genes. Overall, these findings indicate broadly reduced expression of genes involved in intercellular signaling programs required for ongoing kidney morphogenesis in dioxin-exposed kidneys.

The leading-edge genes underlying these development-related processes substantially overlapped with those driving the negative enrichment of the nephrogenic zone signature (Fig. 2I). Nephrons form in the outermost zone of the developing kidney through successive stages of nephron progenitor differentiation into pretubular aggregates, renal vesicles, comma-shaped bodies and S-shaped bodies [35]. Shared leading-edge genes of the nephrogenic zone and morphogenesis included several developmental regulators (*Pax2, Lhx1, Sall1, Robo2, Sfrp2, Hey1*), linking the reduced nephrogenic-zone signature to the broader alteration of kidney developmental signaling [34]. In addition, cell cycle genes (*Top2a, Fbxo5, Pbk*) also contributed to the nephrogenic-zone leading edge, and most individual significantly reduced nephrogenic zone population contained a substantial proportion of cell cycle associated genes (Supplementary file 1; Tab GSEA_E18.5 clusters). Thus, the negative enrichment of the nephrogenic zone signature reflected both altered expression of developmental and cell cycling programs.

The renal stroma is a prominent signaling population in the developing kidney that supports nephron formation, ureteric branching, vascular development and structural organization. As the kidney matures, the relative abundance of stromal cells declines, with the persisting stromal-derived populations comprising fibroblasts, vascular mural cells, and vascular smooth muscle cells, some of which can contribute to fibrosis following injury [36, 37]. In dioxin-exposed kidneys, the renal stromal signature was negatively enriched, with leading-edge genes overlapping with morphogenesis– and cell cycle-related processes (Fig. 2I), but mainly driven by multiple genes coding for extracellular matrix components (*Dcn, Col1a2, Col3a1, Emilin1, Fn1*). Further exploration of stroma-expressed genes previously implicated as regulators of renal morphogenesis through genetic perturbation studies showed coordinated lower expression following dioxin exposure, although only two were individually differentially expressed (Fig. 3D) [36]. Despite these transcriptional changes, PDGFRA/PDGFRB-positive area (Welch’s t-test, p=0.2340) and FN1-positive area (Welch’s t-test, p=0.9185) did not differ between groups, and no alterations in spatial distribution suggestive of fibrotic remodeling were observed (Fig. S2-3). A previous study in C57BL/6N mice reported reduced FN1 staining intensity without altered distribution, alongside alterations of the basal-lamina following dioxin exposure [23]. The absence of comparable protein-level changes may reflect substrain-dependent renal susceptibility [38]. Overall, the coordinated downregulation of stromal genes was not reflected by reduced stromal marker-positive area or altered marker localization, suggesting transcriptional changes within a largely preserved stromal compartment rather than overt stromal loss or fibrosis.

**Figure 3.**
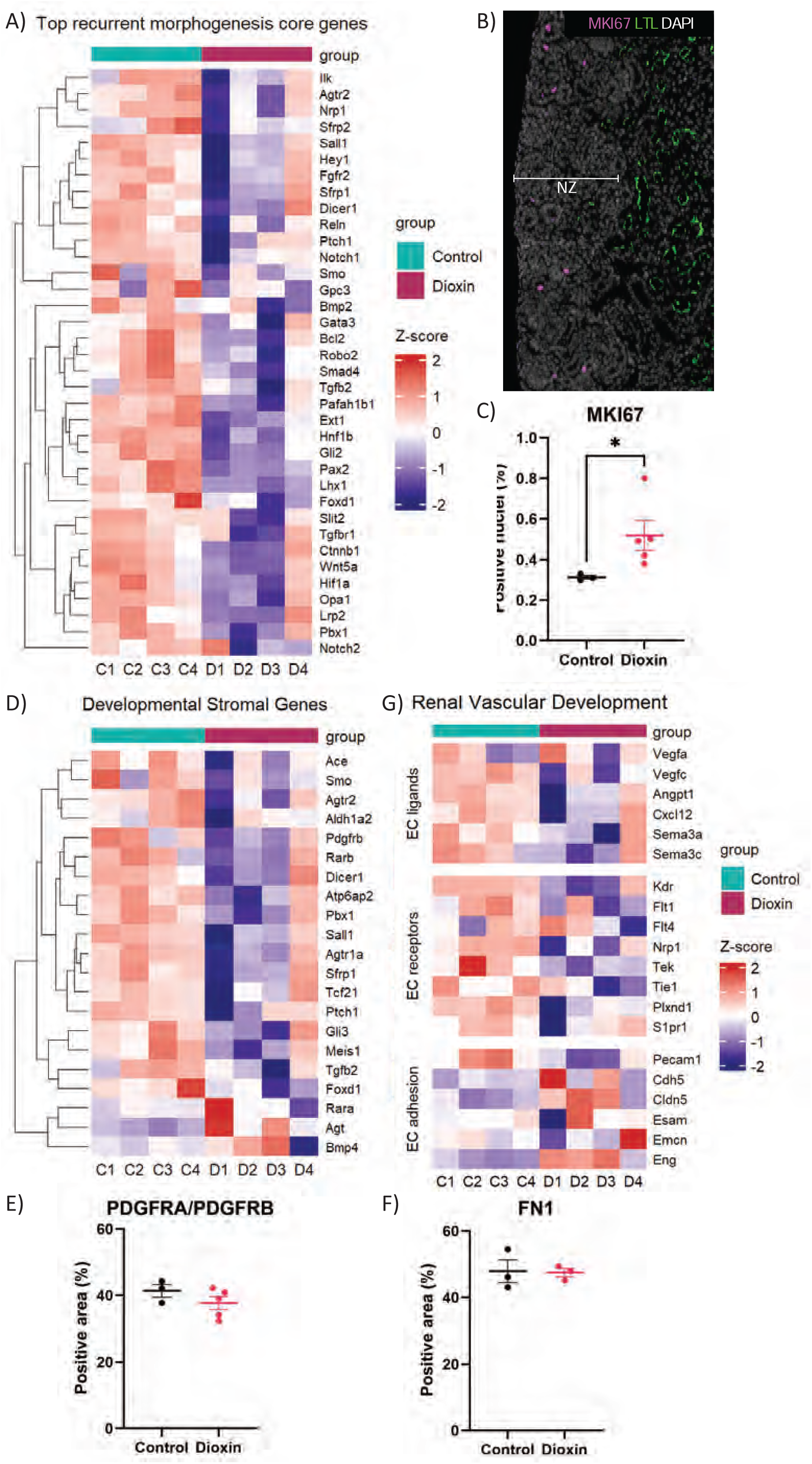
| Dioxin alters developmental programs in the fetal kidney. A) Heatmap of top recurrent morphogenesis core genes. B) Representative MKI67 staining in the nephrogenic zone (NZ). C) Percentage of MKI67-positive nuclei. D) Heatmap of stromal genes linked to kidney developmental defects. E-F) PDGFRA/PDGFRB-positive and FN1-positive area. G) Heatmap of renal vascular-development genes grouped into EC-directed ligands, EC receptors, and EC adhesion molecules. Data are represented as mean ± SEM. p < 0.05.

The renal vasculature is established through a combination of vasculogenesis from endothelial precursors and angiogenic expansion and remodeling of pre-existing vessels [39]. GSEA showed negative enrichment of biological processes related to endothelial development and sprouting angiogenesis following dioxin exposure. Consistently, the vasculature signature was negatively enriched, driven by lower expression of key endothelial genes (*Pecam1, Kdr, Tek, and Fabp4*) [40, 41]. To further examine vascular developmental signaling, selected genes were grouped into endothelial cell (EC)-directed ligands, EC receptors, and EC adhesion molecules, representing signaling input, signal reception, and endothelial organization, respectively [42]. EC-directed ligands and their receptors showed coordinated negative fold changes, whereas adhesion genes displayed mixed changes (Fig. 3E). Together, these findings indicate altered endothelial developmental signaling rather than uniform loss of endothelial identity. Consistent with this, obstruction-associated glomerular alterations have been reported in fetal sheep and non-human primates, while neonatal rats showed changes in both glomeruli and peritubular capillaries [43–45]. The transcriptional changes observed in endothelial signaling may be related to the medullary flattening, given the high density of peritubular capillaries in the developing medulla [46].

### Dioxin exposure dysregulates the cell cycle

During nephrogenesis, cell proliferation supports expansion and maintenance of progenitor populations and formation of nephrons, with cell cycle activity varying across nephron progenitor and differentiating renal cell states [47]. GSEA revealed negative enrichment of 90 cell cycle-related terms involving mitotic progression, chromosome segregation, spindle organization, kinetochore-microtubule attachment, metaphase-to-anaphase transition, and cytokinesis. Recurring leading-edge genes included several core components of the mitotic machinery such as kinetochore-associated genes (*Ndc80*, *Nuf2*, *Ska3*, *Cenpe*), mitotic regulators (*Ccnb1*, *Cdc20*), and all components of the chromosomal passenger complex (*Aurkb*, *Birc5*, Cdca8, *Incenp)* [48–51]. Together, the negative enrichment of these interconnected processes indicates dysregulated activity of the cell cycle machinery.

Given the transcriptional evidence for dysregulation of the cell cycle, we assessed cell cycling by immunofluorescence staining for MKI67. In both control and dioxin-exposed kidneys, high-intensity MKI67-positive nuclei were predominantly located in the nephrogenic zone (Fig. 3B), consistent with ongoing proliferation of nephron progenitors and developing nephron structures, with weaker staining along the outer renal border and few high-intensity positive nuclei in the medulla (Fig S1)[52]. As weaker staining could not be reliably distinguished from background, quantification was restricted to high-intensity MKI67-positive nuclei. The proportion of strongly MKI67-positive nuclei was increased following dioxin exposure (0.31% to 0.52%, Welch’s t-test, p=0.0499; Fig. 3C), despite lower *Mki67* transcript levels (log2FC=−0.87, p<0.05, FDR=0.056) and broad negative enrichment of cell-cycle programs. As MKI67 positivity reflects active cell cycle status rather than proliferation rate itself, these findings suggest altered cell cycle dynamics rather than increased proliferation per se [53]. Overall, the increase in strongly MKI67-positive nuclei despite negative enrichment of cell cycle-related transcriptional programs points to altered cell cycle regulation in hydronephrotic kidneys, although the underlying mechanisms remain unresolved.

### Hydronephrotic kidneys show altered metabolic and xenobiotic-response programs

During nephrogenesis, proliferating nephron progenitors rely predominantly on glycolysis, whereas differentiation is accompanied by a shift toward mitochondrial oxidative metabolism [54]. As the leading-edge genes of the 97 positively enriched metabolic processes included a strong representation of oxidative phosphorylation (OXPHOS) components, we further resolved this signal with GSEA using MitoCarta3.0 MitoPathways [55]. This confirmed positive enrichment of OXPHOS and its respiratory chain complexes, particularly complexes I, IV and V, driven predominantly by nuclear-genome encoded *Nduf*, *Cox*, and *Atp5* gene families (Fig. 4A-B). The enrichment of mitochondrial ribosome and translation pathways were similarly driven by the nuclear-encoded *Mrpl* and *Mrps* families, which code for the proteins that form the mitochondrial ribosome required for translating mitochondrial DNA (mtDNA) encoded transcripts [56]. Notably, no terms related to the mtDNA itself were significantly enriched. Thus, the metabolic dysregulation in the dioxin group predominantly reflected coordinated upregulation of nuclear-encoded mitochondrial OXPHOS and translation machinery, potentially indicating a shift toward a more differentiated metabolic state [54].

**Figure 4.**
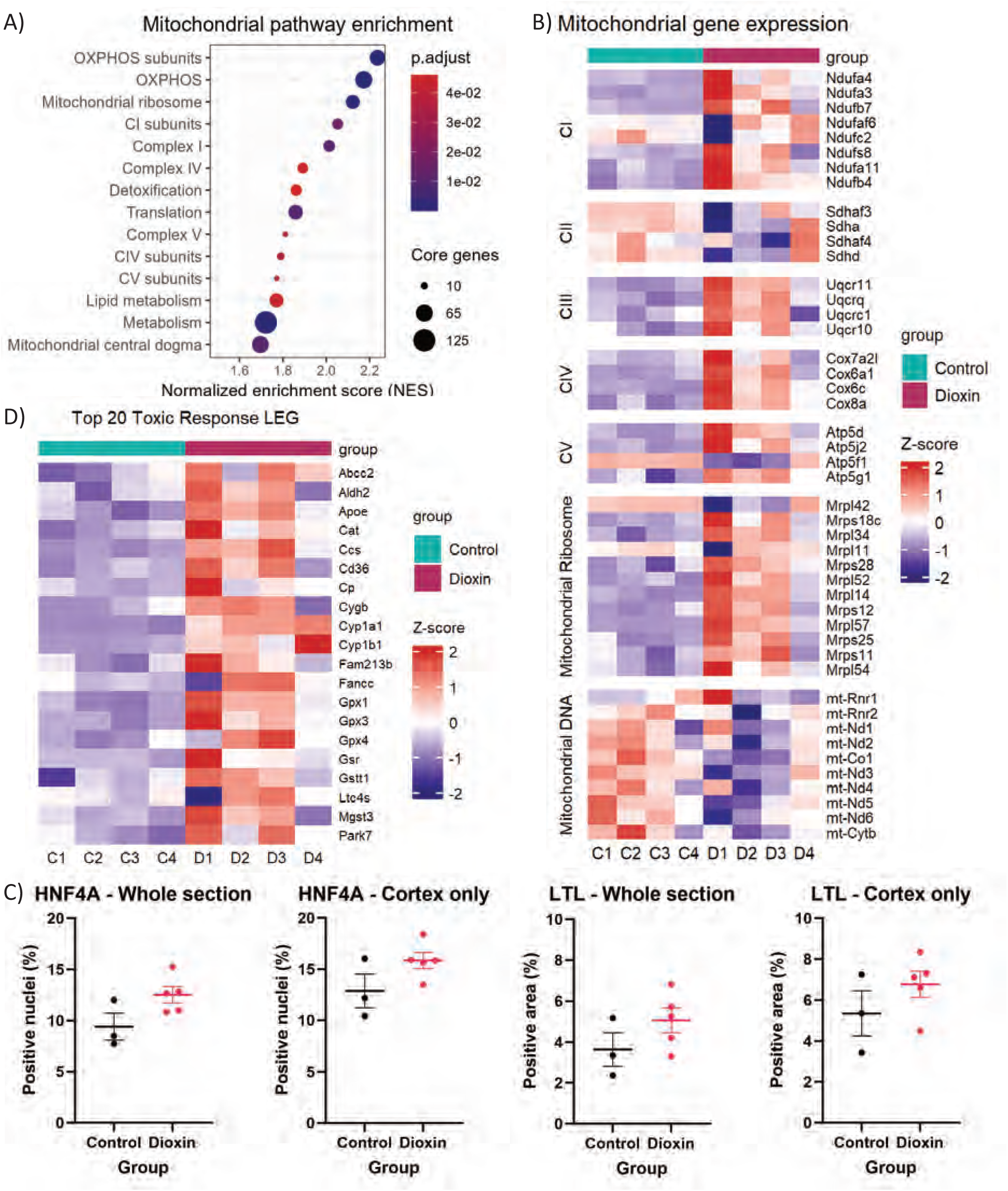
| Dioxin alters mitochondrial and toxic-response programs in the fetal kidney. A) Dot plot of GSEA results for mitochondrial pathways. B) Heatmap of mitochondrial genes grouped by nuclear-encoded respiratory chain complexes I-V and mitochondrial ribosomal genes, and mitochondrial DNA-encoded genes. C) Percentage of HNF4A-positive nuclei and LTL-positive area quantified in whole-kidney sections and cortex separately. D) Heatmap of the top 20 recurrent toxic-response leading-edge genes. Data are presented as mean ± SEM.

The proximal tubule is also a major site of renal xenobiotic metabolism and elimination [57]. Consistent with dioxin-mediated AhR activation, detoxification-related processes were positively enriched and driven by genes involved in Phase I cytochrome P450-mediated xenobiotic metabolism (*Cyp1a1, Cyp1b1*), Phase II glutathione S-transferase-mediated conjugation (*Gstt1, Mgst3*), and glutathione-dependent antioxidant defense (*Gpx1, Gpx3, Gpx4*) [58–60](Fig. 4D). Transcription factor motif enrichment further identified significant enrichment of the canonical AHR:ARNT motif (TF:M00237) among upregulated DEGs (140/412), and both AHR (TF:M00976) and AHR:ARNT (TF:M00237) motifs among downregulated DEGs (49/155 and 64/155, respectively; 86/155 unique genes) [59]. Moreover, toxin-response leading-edge genes most strongly mapped to the complete gene set expressed in the (early) proximal tubule population (34.5%) (Fig S5). Together, these findings support a persistent AhR-associated xenobiotic and antioxidant response in the dioxin-exposed kidney at E18.5, with the proximal tubule likely representing a major site of this response, consistent with its established role in the uptake, metabolism, and excretion of xenobiotics [57].

Metabolic and toxin-response leading-edge genes showed the strongest overlap with the (early) proximal tubule signature (Fig. 2I), which was itself positively enriched and driven by several proximal tubule markers (*Slc34a1, Slc22a6*, and *Aldob*) [61, 62]. The quantification of HNF4A-positive nuclei and LTL-positive area to determine whether this signature reflected an increase in proximal tubule cell abundance, revealed no differences between groups (Welch’s t-test; whole section: HNF4A p = 0.1234, LTL: p = 0.3354; cortex only: HNF4A p = 0.2047, LTL p=0.2308; Fig 5C), consistent with unchanged *Hnf4a* transcript levels (Fig. S4). As HNF4A promotes proximal tubule differentiation [63], the absence of changes in this proximal tubule marker suggests that the enriched proximal tubule signature reflects an altered transcriptional state, potentially related to metabolic and detoxification functions, rather than increased proximal tubule abundance.

### Immune alterations in dioxin group involve innate myeloid-associated programs

Given that inflammation contributes to renal injury following urinary tract obstruction during kidney development, we further explored the immune response in dioxin-exposed kidneys [64]. GSEA identified positive enrichment of a few immune-related processes, including MHC class I and II antigen presentation (*H2-D1, H2-Ab1*) and chemokine-mediated leukocyte recruitment (*Ccl2, Cx3cl1*), together with neutrophil/monocyte-associated (*S100a8, S100a9*) and renal macrophage-associated genes (*C1qa, Apoe*) (Supplementary file 1) [65–67]. In addition, E18.5 immune cell signatures were positively enriched and partly driven by the same myeloid-associated genes. The lack of changes in T-cell or B-cell adaptive immune response suggests that these findings primarily reflect alterations in the innate myeloid-associated immune response in dioxin-exposed kidneys.

## Discussion

In this study, we characterized the morphological and transcriptomic changes associated with dioxin-induced COU. We found that dioxin exposure at E10.5 produced reproducible bilateral hydroureteronephrosis with selective medullary flattening at E18.5. At the molecular level, hydronephrotic kidneys exhibited disruption of developmental and cell cycle pathways, including reduced expression of nephrogenic zone, stromal, and vascular signatures alongside positive enrichment of mitochondrial, proximal tubular, innate immune and toxin-responsive programs. Because these transcriptional alterations were not accompanied by detectable differences in stromal or proximal tubular cell abundance or cortical architecture by immunohistochemistry, they likely reflect disrupted renal developmental trajectories and altered cellular signaling, rather than simply reflecting focal tissue loss or changes in cellular composition. Despite the combined effects of dioxin exposure and obstruction, this reproducible bilateral hydroureteronephrosis phenotype recapitulates several renal features of early human COU, supporting its use as a non-genetic model to study how congenital obstruction perturbs the developing kidney.

A prominent feature of the transcriptional response was the coordinated suppression of genes involved in kidney morphogenesis, cell-cycle, nephrogenic zone, stromal, and vascular gene programs. As kidney development depends on extensive interactions between these compartments, such widespread changes suggest disruption of developmental communication rather than isolated defects [34, 42, 47]. Notably, these transcriptomic alterations occurred without evidence of uniform compartment loss. Although reduced expression of cell cycle programs could reflect impaired nephron progenitor maintenance and nephrogenesis, the concurrent increase in strongly MKI67-positive nuclei points instead to altered cell cycle regulation [68]. Collectively, these findings indicate that congenital obstruction is associated with broad perturbation of renal developmental programs and cellular states within otherwise preserved renal compartments. Together with previous observations that urinary tract obstruction can interfere with nephrogenesis resulting in reduced nephron numbers [8], this suggests that early onset of altered developmental programming may be a central feature of the fetal obstructive response.

During nephrogenesis, proliferative progenitor cells progressively differentiate into mature renal cell types while undergoing a metabolic transition toward increased mitochondrial OXPHOS [54, 69–71]. The dioxin-exposed kidneys showed disruption of these processes, with reduced expression of genes required for developmental and mitotic programs alongside increased expression of nuclear-encoded OXPHOS genes and mitochondrial translation machinery. The enrichment of proximal tubule-associated signatures is compatible with a premature shift toward a more differentiated metabolic state, as mature proximal tubular cells are highly dependent on oxidative metabolism [54]. However, unchanged HNF4A-positive cell abundance and LTL-positive area argue against selective expansion of this compartment and increased numbers of strongly MKI67-positive nuclei indicate that the observed transcriptional changes cannot simply be explained by an increase of mature quiescent cells [53]. Despite the lack of histological evidence for increased proximal tubule abundance, the developmental, cell cycle, and metabolic alterations suggest that obstruction perturbs normal renal developmental trajectories and proliferation-differentiation dynamics.

The response observed in hydronephrotic fetal kidneys differs markedly from that typically reported following obstruction of the mature kidney. Direct comparison is complicated by the substantial overlap between developmental and injury-response pathways, as programs active during kidney development may be reactivated after injury but serve different biological functions [72–74]. In postnatal and adult UUO, injury is commonly associated with proliferative repair responses, developmental gene re-expression, inflammation and fibrosis, together with impaired mitochondrial oxidative metabolism and increased reliance on glycolysis [72, 75–77]. In contrast, hydronephrotic kidneys in the fetal dioxin model exhibited negative enrichment of developmental and cell-cycle programs, enhanced expression of OXPHOS-associated pathways, and no evidence of fibrotic remodeling despite alterations in the innate myeloid-associated immune response. The absence of fibrosis may reflect the relatively short duration of obstruction, although longitudinal studies are needed to determine whether fibrotic remodeling develops at later stages. These differences observed at E18.5 support that obstruction during nephrogenesis does not simply represent mature obstructive injury occurring in a developing kidney, but rather the initial disturbances shaped by the developmental state of the kidney [8]. However, the increase in strongly MKI67-positive nuclei, consistent with an injury-associated proliferative response, contrasts with the negative enrichment of cell cycle programs, suggesting altered cell cycle dynamics or divergent proliferative responses across renal cell populations. Overall, the divergence of the dioxin model from other established models makes it particularly valuable for studying the early developmental consequences of COU.

An important point of convergence with an established feature of obstructive nephropathy in mice was the positive enrichment of innate myeloid-associated immune programs [78]. Inflammatory mediators have likewise been detected in human fetal urine from pregnancies affected by posterior urethral valves [64, 79], supporting inflammation as a shared feature of congenital and postnatal obstructive kidney injury. However, macrophage-associated transcriptional changes should not be interpreted solely as evidence of damaging inflammation, as renal macrophages also contribute to developmental remodeling, clearance of cellular debris, and tissue repair [80]. Consequently, the immune signature observed here may reflect a combination of inflammatory and adaptive tissue-remodeling responses to congenital obstruction.

Early fetal COU in humans can present with medullary flattening while overall renal architecture remains largely preserved, preceding overt tubular injury and tubulointerstitial fibrosis [6, 81, 82]. Consistent with early fetal COU, dioxin-exposed kidneys at E18.5 display features of early obstruction, including medullary flattening, preservation of overall renal structure, and a lack of histological evidence for tubular injury or fibrosis. The morphological changes have been reported to coincide with the onset of dioxin-induced ureteral obstruction around E15.5 [20, 23]. Nevertheless, the persistent AhR-responsive xenobiotic signature observed at E18.5 indicates that the renal phenotype reflects both congenital obstruction and continuing molecular consequences of prenatal dioxin exposure.

Despite the combined contribution of dioxin-exposure and obstruction, the dioxin model is one of the few non-genetic models that reproducibly induces bilateral hydroureteronephrosis in utero while recapitulating renal features relevant to early human COU. In addition, other compound-induced experimental kidney injury models, including high-dose folic acid nephropathy, have proven to provide insight into downstream pathogenic mechanisms without mirroring the human etiology [83]. Dioxin-induced hydroureteronephrosis therefore offers a valuable platform to investigate how fetal obstructive uropathy perturbs kidney development. At E18.5, informative readouts include pelvic dilatation, medullary flattening, altered developmental programs and accompanying metabolic and inflammatory responses. These features provide opportunities to test whether genetic or environmental modifiers influence susceptibility to obstruction-induced disruption of renal development. Furthermore, because animals exposed to dioxin in utero survive postnatally, this model enables longitudinal studies to determine whether early developmental disturbances have lasting consequences for nephron endowment and kidney function.

Several limitations should be considered when interpreting these findings. Bulk RNA sequencing cannot distinguish changes in cell abundance from changes in transcriptional state. Single-cell or spatial approaches will therefore be needed to attribute the dysregulated biological processes to specific cell types and determine whether they reflect altered cellular composition or cell-intrinsic transcriptional changes. In addition, substantial interindividual variability in the transcriptional response to dioxin was observed. Despite gross hydroureteronephrosis comparable to other dioxin-exposed kidneys, sample D4 clustered transcriptionally closer to controls. Although it retained a toxin-associated response, particularly *Cyp1a1* induction, its inclusion attenuated rather than drove the transcriptional differences between groups. This variability could reflect differences in obstruction severity, litter-associated factors, or individual biological responses to dioxin exposure. Furthermore, histology and RNA sequencing were performed respectively on the left and the right kidney, thus precluding direct morphological and transcriptional comparisons. This is further complicated by the fact that right-sided lesions have previously been reported to be more severe [84]. The analysis was limited to a single late-gestation snapshot, thereby impeding the determination of whether the E18.5 phenotype is transient or results in lasting developmental abnormalities. Although mouse nephrogenesis continues until postnatal day 3–4 [34, 85] and dioxin-associated hydronephrosis can partially regress after birth [84], disruption during late nephrogenesis could reduce final nephron endowment and renal resilience [86]. Postnatal assessment of nephron number, renal structure, and function is therefore needed to determine the long-term consequences of these prenatal changes. Lastly, comparison with additional developmental obstruction models will also be required to distinguish the response to obstruction from dioxin-specific effects.

To our knowledge, this is the first characterization of the broad transcriptional response of the developing mouse kidney to congenital urinary tract obstruction at E18.5. Overall, dioxin-induced hydroureteronephrosis captures an early developmental response to fetal urinary tract obstruction characterized by medullary flattening in a relatively preserved kidney, broad disruption of developmental signaling, altered cell cycle regulation, a distinct oxidative metabolic response, and innate myeloid-associated immune response changes. Its divergence from classical adult UUO emphasizes that obstruction occurring during nephrogenesis represents a developmental insult whose consequences depend on the developmental state of the kidney. The model therefore provides a platform to investigate how prenatal obstruction alters ongoing renal development and whether these disturbances ultimately affect nephron endowment and long-term kidney health.

## Materials and methods

### Preparation of dioxin and control stock solution in corn oil

Dioxin (#DRE-GA09011171TO, LGC standards), supplied in toluene, was transferred into corn oil (#C8267, Sigma-Aldrich) using a gravimetric approach. 1 mL of the dioxin-toluene solution was added to a pre-weighed glass bottle, followed by 1 mL corn oil, and the bottle was weighed after each step. The toluene was then removed by evaporation at room temperature in a fume hood under continuous stirring, monitoring weight loss until it matched the mass of the glass bottle and oil, confirming complete evaporation. Corn oil was then added to reach a final concentration of 1.5 µg dioxin/mL. Control solutions were prepared identically, using an equivalent volume of toluene without dioxin, to control for trace toluene residue. The solutions were stored in the dark at 4°C.

### Animals and treatments

All mice were maintained in individually ventilated cages at 20-22°C on a 12-h light/dark cycle, with unrestricted access to water and standard RM3 chow (SDS, Essex, United Kingdom). All animal procedures were approved by the Animal Welfare Body Leiden and conducted in accordance with the Dutch Experiments on Animals Act and EU Directive 2010/63/EU. For the experiment male and female Cdh12^ΔSCON/+^ mice were used. The generation of the Cdh12^ΔSCON^ mouse line (C57BL/6JLumc-Cdh12em1.1Lumc) was previously described [87]. Briefly, the line was established through breeding to obtain germline transmission of the ΔSCON allele and segregation of the Cre transgene, followed by crossing with C57BL/6J mice to generate Cdh12^ΔSCON/+^ animals carrying one ΔSCON allele and one wildtype Cdh12 allele.

Timed matings were set up overnight with male and female Cdh12^ΔSCON/+^ mice. Although these intercrosses generated both wildtype and mutant fetuses, the present study focuses on the characterization of the dioxin model in wildtype fetuses. Females likely to be pregnant based on the presence of a copulatory plug and/or post-mating weight gain received dioxin by oral gavage at E10.5 (15 µg/kg, dosed by body weight on the day of administration) or an equivalent volume of control solution. This dose was selected as the lowest previously shown to induce 100% penetrance of hydroureteronephrosis [88]. Female body weight was assessed at day 0.5, 7.5, 10.5, 11.5, 12.5, 13.5, 14.5, 17.5, and 18.5 post mating from which the percentage change in body weight was calculated relative to day 0.5 post mating. At E18.5, dams were euthanized by CO₂ inhalation, after which fetuses were isolated and decapitated. Per litter the total number of implantation sites, the number of live fetuses (no dead fetuses were found in any group) and resorptions were assessed. Fetuses were classified as alive based on their response to gentle stimulation, while resorptions were identified by signs of tissue degeneration. Although these litters contained wildtype, Cdh12^ΔSCON/+^ and Cdh12^ΔSCON/ΔSCON^ fetuses, previous studies demonstrated that these genotypes are viable and occur at the expected Mendelian ratios [87]. Hence, genotype was not expected to influence litter outcomes.

Fetuses were maintained in ice-cold PBS until dissection and examined under a stereomicroscope for hydroureteronephrosis. Right kidneys were snap-frozen in liquid nitrogen and stored at –80°C for RNA extraction, whereas left kidneys were fixed overnight in 4% PFA and subsequently paraffin-embedded for histological analysis. Genomic DNA was isolated from tail biopsies by alkaline lysis (25 mM NaOH, 0.2 mM EDTA) for 30 min at 95°C, followed by the addition of an equal volume of neutralization buffer (40 mM Tris-HCl) after cooling. Genotyping was subsequently performed by PCR as previously described [87].

### Immunohistochemistry

To assess overall kidney morphology, 4-μm-thick coronal sections of paraffin-embedded kidneys were deparaffinized, rehydrated, and stained with H&E. Slides were digitized using a PANNORAMIC P480 slide scanner (3DHISTECH) and analyzed using QuPath version 0.7.0 [89]. For each kidney, the section with the largest craniocaudal length was selected for morphometric analysis. Kidney length was measured along the craniocaudal axis, with cortical thickness at the cranial and caudal poles defined as the distance from the pole to the corticomedullary junction. At the interpolar midpoint, parenchymal thickness was measured from the capsule to the pelvis, and cortical thickness from the capsule to the corticomedullary junction. Medullary thickness was calculated by subtracting cortical from parenchymal thickness. Except for craniocaudal kidney length, all measurements were expressed as a percentage of craniocaudal length to account for differences in kidney size.

For immunofluorescence staining, deparaffinized and rehydrated sections underwent heat-induced antigen retrieval (20 min in pressure cooker) using citrate buffer (pH 6) or Tris-EDTA buffer (pH 9). Sections were subsequently blocked using the M.O.M. Basic Kit (#BMK-2202, Vector Laboratories) and incubated with primary antibodies, followed by appropriate fluorescent secondary antibodies. Antibodies and staining conditions are listed in Supplementary Table S1. Slides were digitized using a Axioscan 7 slide scanner (ZEISS), and images were quantified using QuPath version 0.7.0 [89].

### 3’ end bulk RNA sequencing

Total RNA was isolated and purified from whole kidney tissue lysates collected from E18.5 wildtype fetuses using the RNeasy Mini Kit (#74104, Qiagen). RNA quality was assessed on the Agilent 2100 Bioanalyzer (#G2939BA, Agilent Technologies) using the RNA 6000 Nano Kit (#5067-1511, Agilent Technologies) before proceeding to library preparation. Four kidneys per group were analyzed, each derived from a distinct litter (n=4). The cDNA library generation and Illumina sequencing were performed by the Leiden Genome Technology Center library preparation as previously described [90].

Raw sequencing data was processed in Galaxy [91] as follows: adapter sequences were trimmed from R1 and R2 using Cutadapt, followed by trimming of R2 to retain the 8-bp unique molecular identifier (UMI) and linkage of the UMI to the corresponding cDNA-containing R1 read [92, 93]. Next, reads shorter than 30 bp and poly-G tails were removed using fastp [94]. UMIs were extracted using UMI-tools and ribosomal RNA-derived reads were removed using SortMeRNA [95, 96]. The remaining reads were aligned to the mouse reference genome (mm10) using STAR and deduplicated based on genomic position and UMI using UMI-tools [95, 97]. Gene count matrices were subsequently generated using featureCounts with the GENCODE Release M10 (GRCm38.p4) mouse annotation [98]. Data were further analyzed in Rstudio using R version 4.5.0. DEGs were identified using DESeq2, applying thresholds of absolute fold change ≥1.5, *p* <0.05, and a non-missing adjusted *p* value. GO enrichment analysis was performed separately for up– and downregulated DEGs using clusterProfiler, with gene set sizes restricted to 10-500 genes. All GSEA analyses were performed using the complete gene list ranked in descending order by −log10(*p* value) × sign(log2 fold change). For both GO and GSEA terms with an FDR-adjusted p-value < 0.05 were considered significantly enriched. GSEA was conducted using clusterProfiler with curated mouse biological process gene sets, with gene set sizes restricted to 10-500 genes [99] and mitochondrial pathways from MitoCarta3.0 [55]. GSEA of cell population signatures was performed using up to 100 cluster-marker genes per cluster from two E18.5 mouse kidney single-cell RNA-sequencing datasets [32, 33], selected based on log2 fold change and *p* <0.05, with marker specificity additionally considered for one dataset. Based on these markers, corresponding clusters across datasets were matched and grouped into cell populations. Their marker genes were combined to generate cross-study consensus signatures, which were analyzed alongside dataset-specific signatures. AhR motif enrichment was assessed separately for up– and downregulated DEGs using gprofiler2, restricted to the transcription factor database.

## Statistical analysis

For pregnancy outcomes and immunofluorescence statistical analyses were performed using SPSS version 29.0 and GraphPad Prism version 10.6.1, respectively. Continuous data were analyzed using three-way ANOVA or an unpaired two-tailed Student’s t-test, as appropriate. Proportional data were analyzed using the generalized linear model (GLM) with a binomial distribution and logit link. However, near-complete separation between groups prevented reliable model estimation for hydroureteronephrosis, and this outcome was therefore not formally tested using binomial GLM. Immunohistochemical measurements were compared using Welch’s t-test. Statistical significance was set at p<0.05.

## Abbreviations

AhR: Aryl hydrocarbon receptor
BP: Biological process
COU: Congenital obstructive uropathy
DEG: Differentially expressed gene
Dioxin: 2,3,7,8-tetrachlorodibenzo-*p*-dioxin
E: Embryonic day
EC: Endothelial cell
GO: Gene ontology
GSEA: Gene set enrichment analysis
H&E: Hematoxylin and eosin
mtDNA: Mitochondrial DNA
OXPHOS: Oxidative phosphorylation
PCA: Principal component analysis
UUO: Unilateral ureteral obstruction

## Acknowledgements

This study was funded by the Dutch Kidney Foundation through the ArtDECO consortium (no. 20OC002). M.H. and J.M. are a member of the NL Research consortium Kidnie, which is supported by the Dutch Kidney Foundation, and J.M. is a member of the European Reference Network for Rare Kidney Diseases (ERKNet).

## Footnotes

### Author Contributions

M.H., P.H., and J.M. conceived and designed the study. M.H. and R.L. performed the mice handling.

M.H. performed the study and the analysis. K.S. supported with the RNA bulk sequencing analysis.

M.H. wrote the manuscript. M.H., P.H., and J.M., revised and edited the manuscript. All authors reviewed the manuscript.

## Ethics declarations

### Ethical approval

The animal care and experimental procedures were approved by the Animal Welfare Body Leiden and are in accordance with the Dutch Experiments on Animals Act and EU Directive 2010/63/EU.

### Data and resource availability

Data and code supporting this study are available from the corresponding author upon reasonable request. All relevant data and details of resources can be found within the article and its supplementary information.

## Supplementary information

Below is the link to the electronic supplementary information. Supplementary file1

