## Supplementary material for "Morphological and transcriptomic renal characteristics of dioxin-induced fetal obstructive uropathy": Sup Figures

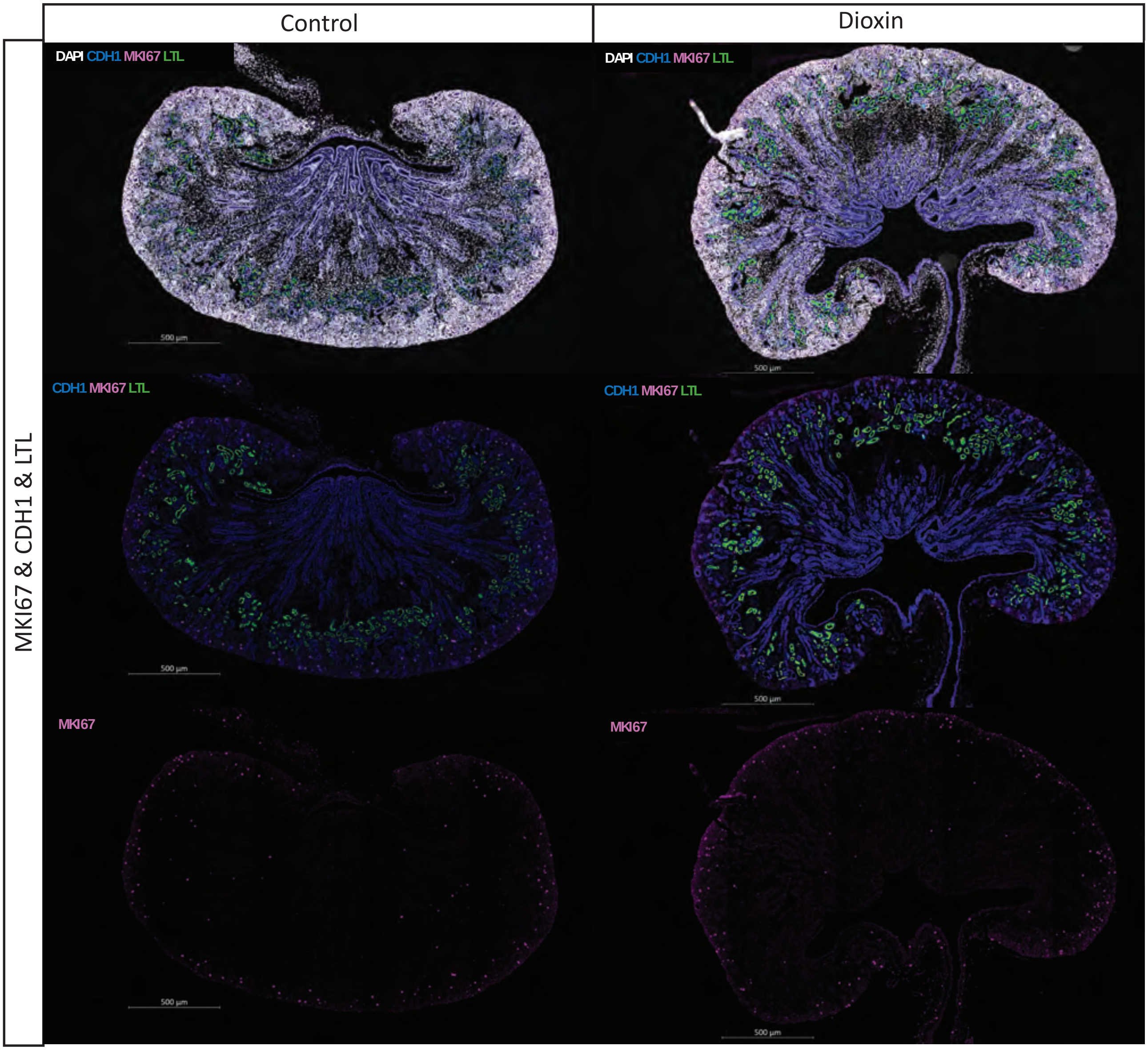


**Figure S1** | Representative immunofluorescence co-staining for MKI67, CDH1, and LTL in control and dioxin-exposed E18.5 kidneys.

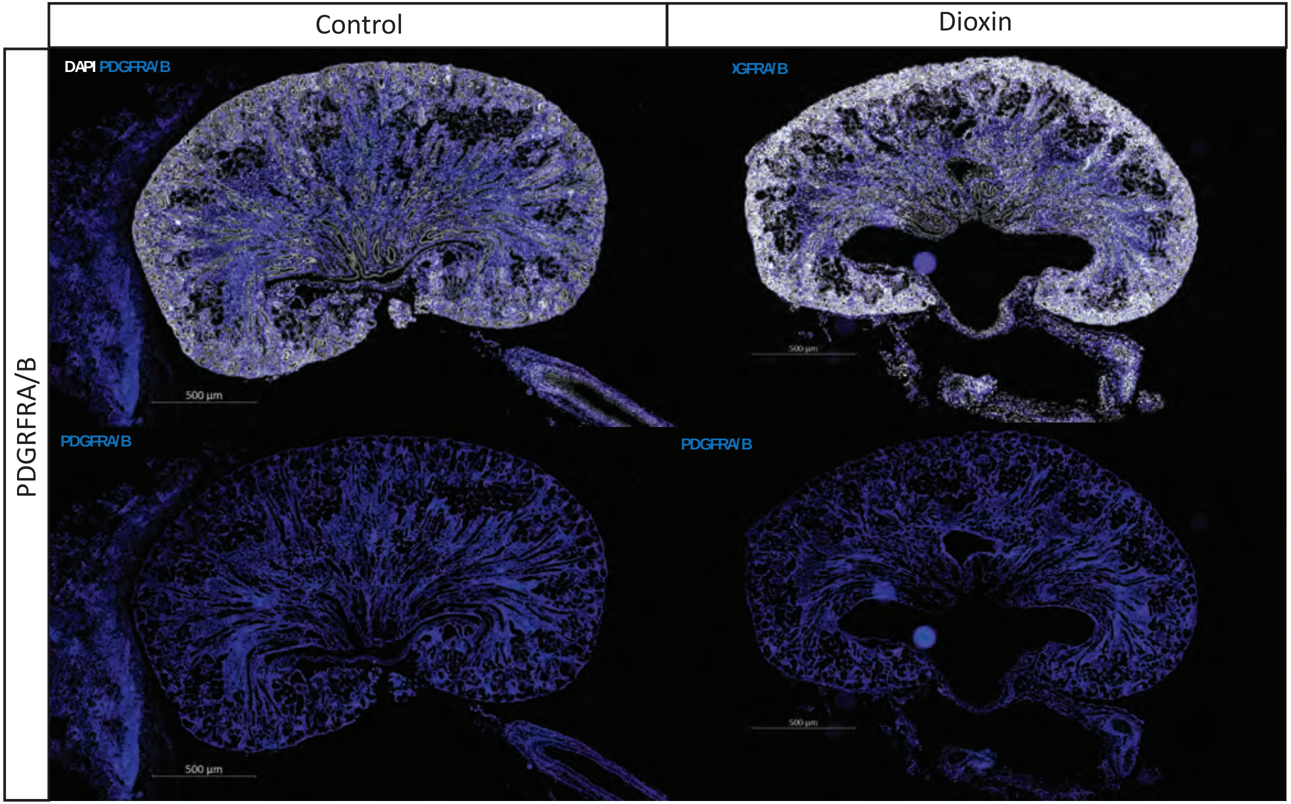


**Figure S2** | Representative immunofluorescence staining for PDGFRA/B in control and dioxin-exposed E18.5 kidneys.

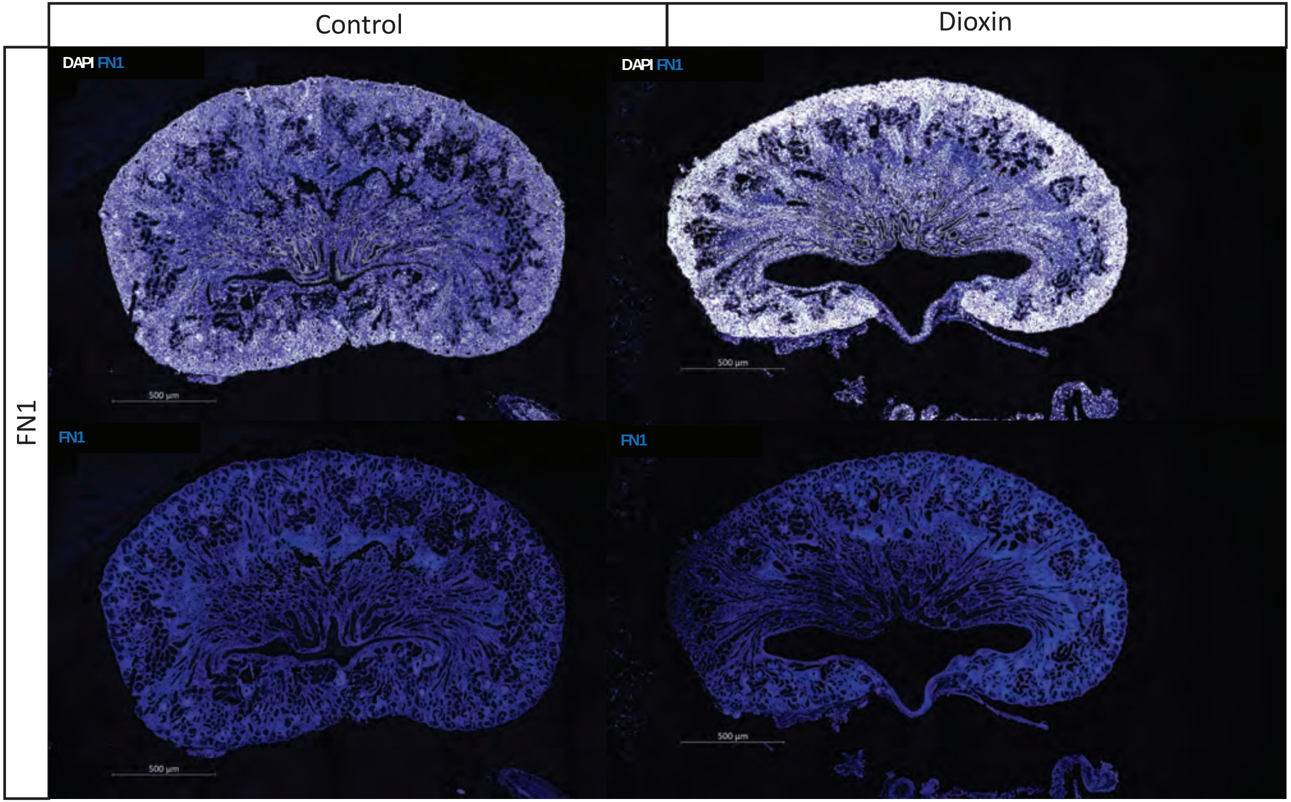


**Figure S3** | Representative immunofluorescence staining for FN1 in control and dioxin-exposed E18.5 kidneys.

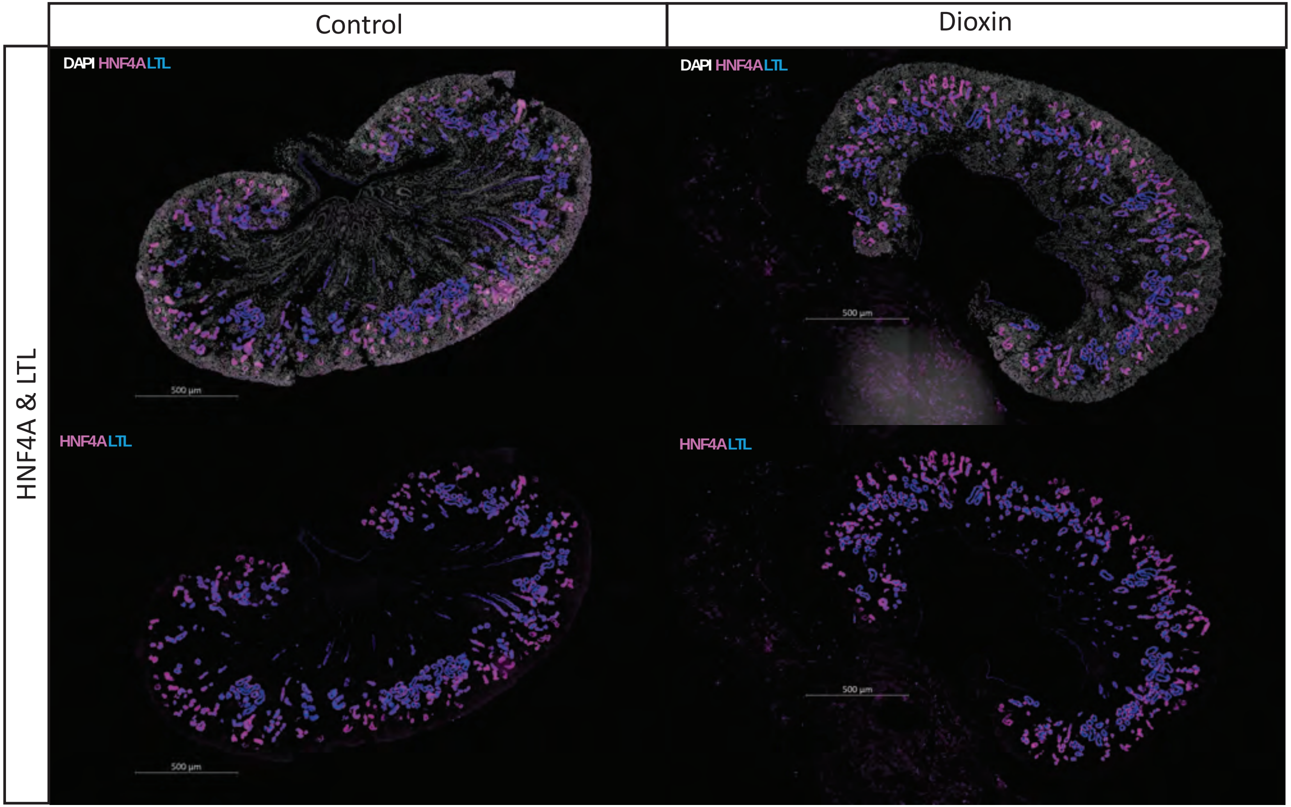


**Figure S4** | Representative immunofluorescence co-staining for HNF4A and LTL control and dioxin-exposed E18.5 kidneys.

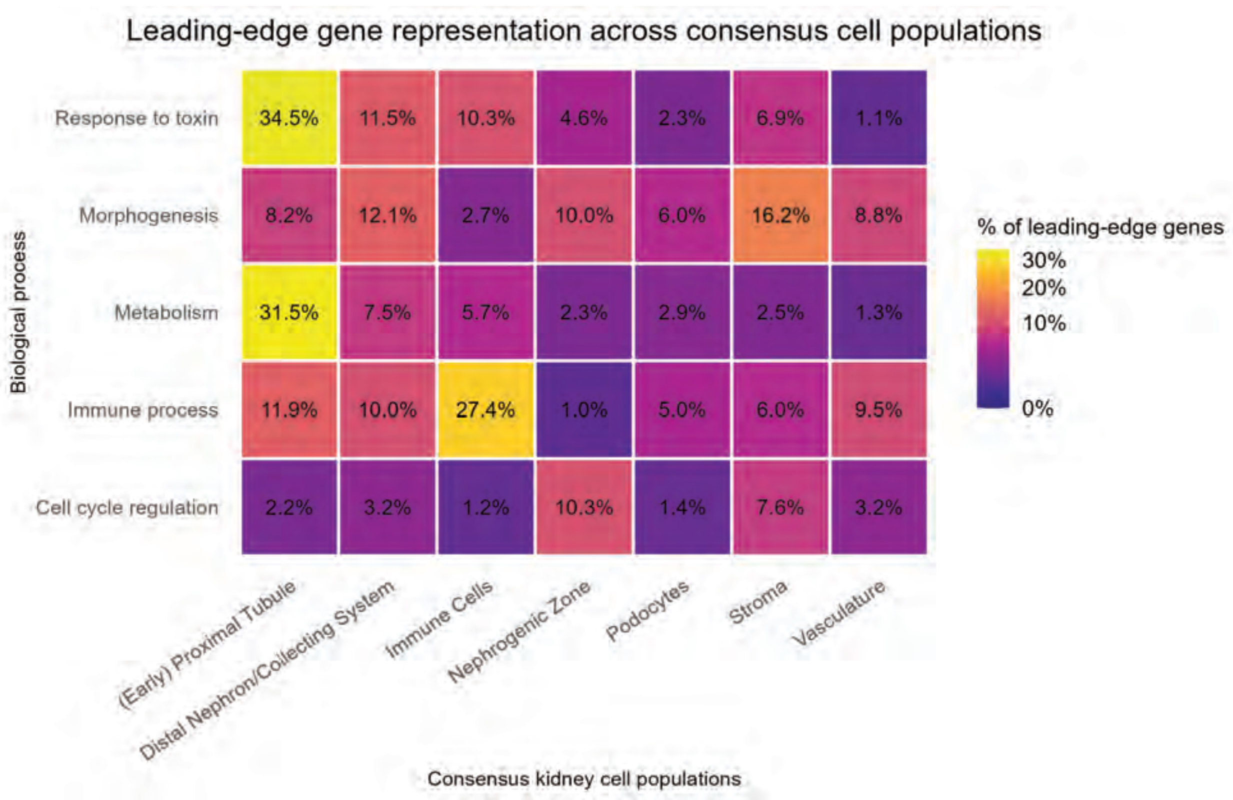


**Figure S5** | Heatmap showing the percentage overlap between leading-edge genes from major enriched biological processes and the complete gene set of consensus kidney cell-population signatures
